# Response-optimised training improves learning of a complex motor task and closely related motor tasks

**DOI:** 10.1101/2025.09.12.675962

**Authors:** Nisha Maria Prabhu, Martin Matke, Nico Lehmann, Norman Aye, Gabriel Ziegler, Marco Taubert

## Abstract

Regular physical exercise is essential for promoting healthy aging and longevity. In older adults with varying physical and cognitive decline, optimising exercise interventions is crucial to maximise benefits. A promising approach to achieve this goal is by adjusting task demands to individual abilities in turn preventing over- or underloading their abilities. In the field of motor learning, it is currently unclear whether such an optimised training improves not only performance on the trained task but also transfers to untrained motor and cognitive tasks.

We conducted a randomised, single-blinded, 6-week dynamic balance training (DBT) with healthy older adults (n=30). Training was tailored to individual balance ability. Participants were assigned to either suboptimal (high or low difficulty) or optimal (moderate difficulty) training groups. Transfer effects were assessed via cognitive tasks (memory and executive) and motor tasks (untrained DBT variations and other balance tasks) measured pre-, mid- and post-intervention. Multivariate longitudinal statistical analysis showed higher performance gains in the optimal training group in three out of six sessions compared to the suboptimal groups, especially under testing conditions with high task demands. The optimal group also showed greater improvements in near motor transfer tasks mid- and post-intervention, while no significant differences were observed in the cognitive tasks. Within-group DBT learning positively correlated with transfer gains, highlighting the role of training response in achieving transfer. In conclusion, optimised task difficulty in balance training enhances both task-specific performance and related motor skills, supporting the use of personalised interventions to maintain function and independence in older adults.

**Highlights:**

- Optimisation of training difficulty based on individuals abilities enables early learning gains.
- Moderate difficulty training may be optimal for improvements in task-specific and untrained near motor transfer tasks.
- Response to training is a vital predictor of transfer potential.

## 1. Introduction

Combating age-related deterioration in quality of life is a growing concern especially with an ever-increasing life expectancy. As older adults experience physical and cognitive decline, their ability to meet the demands of everyday life is affected. Leading an active lifestyle through regular physical exercise can counteract these age-related effects (Boa Sorte Silva et al., 2024; Garatachea et al., 2015). However, during training their ability to keep up with task demands can be limited and vary for every individual (Bootsma et al., 2021). An efficient training plan would ideally balance out the cost of training (time and effort) with not just the task-specific but also transferable improvements. In this regard, motor training that incorporates an aspect of skill learning has the ability to also benefit cognitive health (Tomporowski and Pesce, 2019). Here, additionally optimising task demands may help maximise training gains while avoiding excessive physical and cognitive strain.

In the motor domain, the challenge point framework (CPF) proposes that learning is enhanced when the nominal difficulty of a task is matched to the individual’s skill level and practice conditions (Guadagnoli and Lee, 2004). Too little or too much challenge can hinder learning, therefore, different ability levels require individually tailored task demands (functional task difficulty). Maintaining an optimal challenge point by progressively adapting task difficulty paced to the individuals abilities further supports information processing and motor learning (Hodges and Lohse, 2022). This theoretical framework bears similarities with neuroscientific theories that suggest maintenance of adequate challenge drives behavioural flexibility (Supply-demand mismatch; Lövdén et al., 2010).

Several studies have applied CPF to motor learning tasks such as mirror star tracing (Bootsma et al., 2021, 2020, 2018), postural control (Akizuki and Ohashi, 2015), disc propulsion (Sanli and Lee, 2015), dart-throwing task (Ong et al., 2015), etc. These studies showed that while younger adults benefited from moderate to high nominal difficulty training, healthy older adults profited from moderate to low difficulty levels. However, most studies failed to confirm predictions of the CPF and/or to identify a consistent optimal challenge point that applies across subjects, likely due to lack of individualisation of task demands. These studies applied fixed nominal difficulty levels without accounting for varying individual skill levels (functional task difficulty), potentially contributing to inconclusive outcomes.

When processing demands of distinct tasks overlap, improvements in task-specific performance may underlie transfer (Rogge et al., 2017; Williams and Hodges, 2023). In particular, motor learning relies on cognitive processing (information processing and storage) in order to successfully adapt to challenges (Guadagnoli and Lee, 2004; Tomporowski and Pesce, 2019). Therefore, motor learning can create a potential for transfer, if the trained and the untrained tasks share cognitive or motor components (Lövdén et al., 2010; Taatgen, 2021).

As a result, in this study, we applied CPF principles to dynamic balance training (DBT) in older adults (Sehm et al., 2014), by adjusting the nominal task difficulty to match individual abilities. This was achieved by adding weights under the balance board (making the balance board heavier or lighter) across sessions to maintain the assigned functional task difficulty level. Using this approach, it was possible to manipulate balance board sensitivity and create high (overload), low (underload) and moderate (optimal) difficulty training conditions based on pre-session DBT performance assessments. We examined how these individualised training loads (response-optimised training) affected DBT learning and transfer to motor and cognitive tasks. Transfer was assessed using a comprehensive test battery, comprising of untrained DBT variations (near motor transfer), novel balance task on a static platform (far motor transfer), and cognitive tasks targeting overlapping domains (executive and memory). Here, we target overlapping cognitive domains with neural correlates corresponding with structural regions previously shown to undergo plastic changes after long-term DBT learning (Sehm et al., 2014; Strobach and Karbach, 2016; Taubert et al., 2010). We hypothesised that optimal challenge would yield greater retention and transfer, in line with CPF predictions (Guadagnoli and Lee, 2004).

## 2. Methods

### 2.1. Ethics statement and study design

This study was approved by the local ethics committee of Otto-von-Guericke University, Magdeburg [196/19] and conducted in accordance with the declaration of Helsinki. All subjects provided written informed consent prior to their participation in the experiment and received financial compensation for their participation.

Through this randomized controlled, single-blinded study, we investigated the effect of response-optimised training on learning of a DBT (Wulf et al., 2003) and its associated transfer to untrained motor and cognitive tasks. We examined this effect on individuals between the ages of 60-80yrs of age (n=30, 67.5 ± 4.79y, 18 biological females). Highly skilled participants like slackliners or those with prior DBT experience were excluded. All participants were required to fill-in an activity questionnaire (Huy, 2011) in order to evaluate their general level of physical activity. Additionally, the Montreal Cognitive assessment (MoCA; Nasreddine *et al*., 2005) was conducted to rule out cognitive impairments.

After granting their written informed consent, participants were randomly assigned to either of the three experimental groups, viz., high, moderate or low difficulty training groups (n = 10 per group). The entire training duration lasted a total of 6 weeks consisting of one training session per week (TS1-TS6). Each training session began with an estimation of the participants current ability on the DBT using a balance skill assessment procedure described below. Based on their measured current ability, participants in the high difficulty group received a consistently higher functional difficulty training whereas participants from the low difficulty group received a training which was significantly easier (low functional difficulty). Similarly, the optimal group received a moderate level of task difficulty training (hypothesized as optimal training). Pre- and post-tests were conducted approximately 24 hours before and after the training period respectively. An additional intermediate testing session was conducted immediately preceding the 4^th^ training session (TS4) in order to capture early transfer effects as a result of three training sessions (Fig. 1). The testing sessions (pre, TS4/inter, post) included the DBT balance skill assessment, a variety of motor and cognitive transfer tasks.

**Figure 1.**
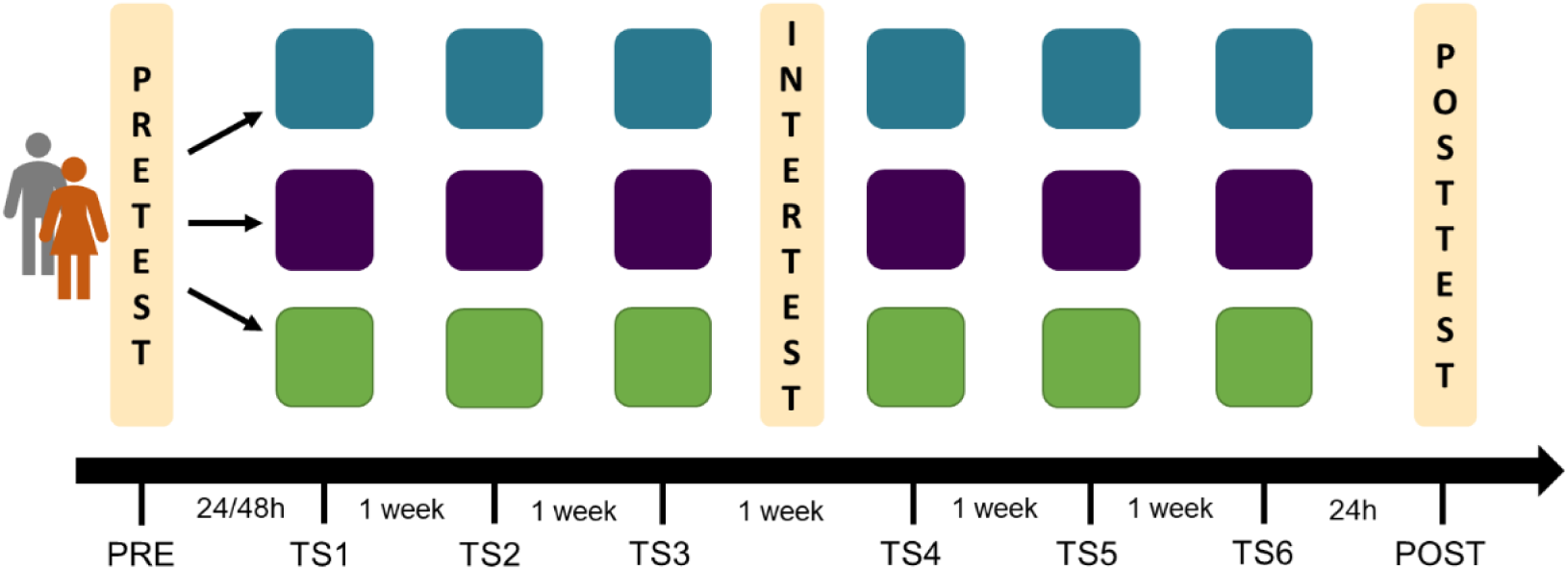
Experimental design: Participants were trained on the DBT over 6 weeks with one practice session every week, the training differed based on their group assignment. Each session included 15 trials lasting 30 s each, interspersed with a rest period of 90 s. Participants also performed a battery of motor and cognitive transfer tests before (pre), midway (inter) and after (post) the training duration.

### 2.2. Complex balance task (DBT)

The motor learning paradigm in our study included a whole-body dynamic balance task known as stabilometer (stability platform, Model 16030, Lafayette Instruments, Lafayette, IN, USA), a balance platform that tilts in a see-saw like manner with a maximum deviation of 26 degrees on each side. Each training session on the stabilometer comprised of 15 trials lasting approximately 30 seconds each trial (Sehm et al., 2014; Taubert et al., 2010). The task as explained to the participant was to maintain the platform in a horizontal position, i.e., parallel to the floor. All participants were asked to stay within a target deviation of 0 - 3° to the right or left from the horizontal axis for as long as possible during each 30s trial. In order to successfully execute this task, the participant is required to position the body’s centre-of-pressure vertically above the boards’ axis of rotation. Participants received a rest period of 90s between each trial. At the end of each trial the participants received feedback about their performance in the form of time in balance (TIB), i.e., seconds spent within the 0 - 3° target window. No directions or instructions regarding task execution strategies were provided apart from the necessary safety guidelines and TIB feedback. Instead, the participants were granted the freedom to explore their own strategies in order to improve performance over the 6 training sessions (Discovery learning approach: Orrell et al., 2006; Wulf et al., 2003). Each training session lasted approximately 45-50 mins each day.

#### DBT skill assessment

This assessment comprised of 6 trials lasting 20secs each conducted before every training session, whereas at pre- & post-test this assessment was repeated twice (6 trials *2). These assessment trials were performed in a pseudo-randomised order with each trial assigned a different level of task difficulty ranging from level 0 (highest difficulty) to level 5 (lowest difficulty), i.e., each level meant adding a metal plate weighing 5kgs each under the board. Adding these weighted plates to the bottom of the board made the board heavier hence less sensitive (more stable) making the task easier. This allowed us to test the participants’ DBT skill on an entire spectrum of nominal task difficulties enabling a prediction of their current skill. Here, based on their performance on each difficulty level, a simple linear model was used to estimate the task difficulty level needed to reach a target time threshold (depending on group assignment). Performance feedback regarding TIB was not provided during the assessment. After every trial the participants received a break of 90 seconds, during which they were asked to perform a short washout task (10m tandem walk).

#### Training procedure

Based on the DBT skill assessment, every training session was designed to start at an individualised training load or functional task difficulty (level 0 – level 7 of nominal task difficulty). Each training session comprised of 15 trials (30 seconds for each trial). Progression of training meant changing the nominal task difficulty after every block of 3 trials (5 blocks in TS1 and TS2) or 5 trials (3 blocks in TS3-TS6) based on their DBT performance, average TIB recorded during these blocks (Fig. 2 & 3). Performance feedback regarding TIB was provided after every trial (CPF: knowledge of results). The threshold for progression and the training load for all the three groups depended on the group assignment, i.e., the threshold for high difficulty group: 3-6 secs; the moderate difficulty group: 12-15 secs; the low difficulty group: 23-26 secs.

**Figure 2.**
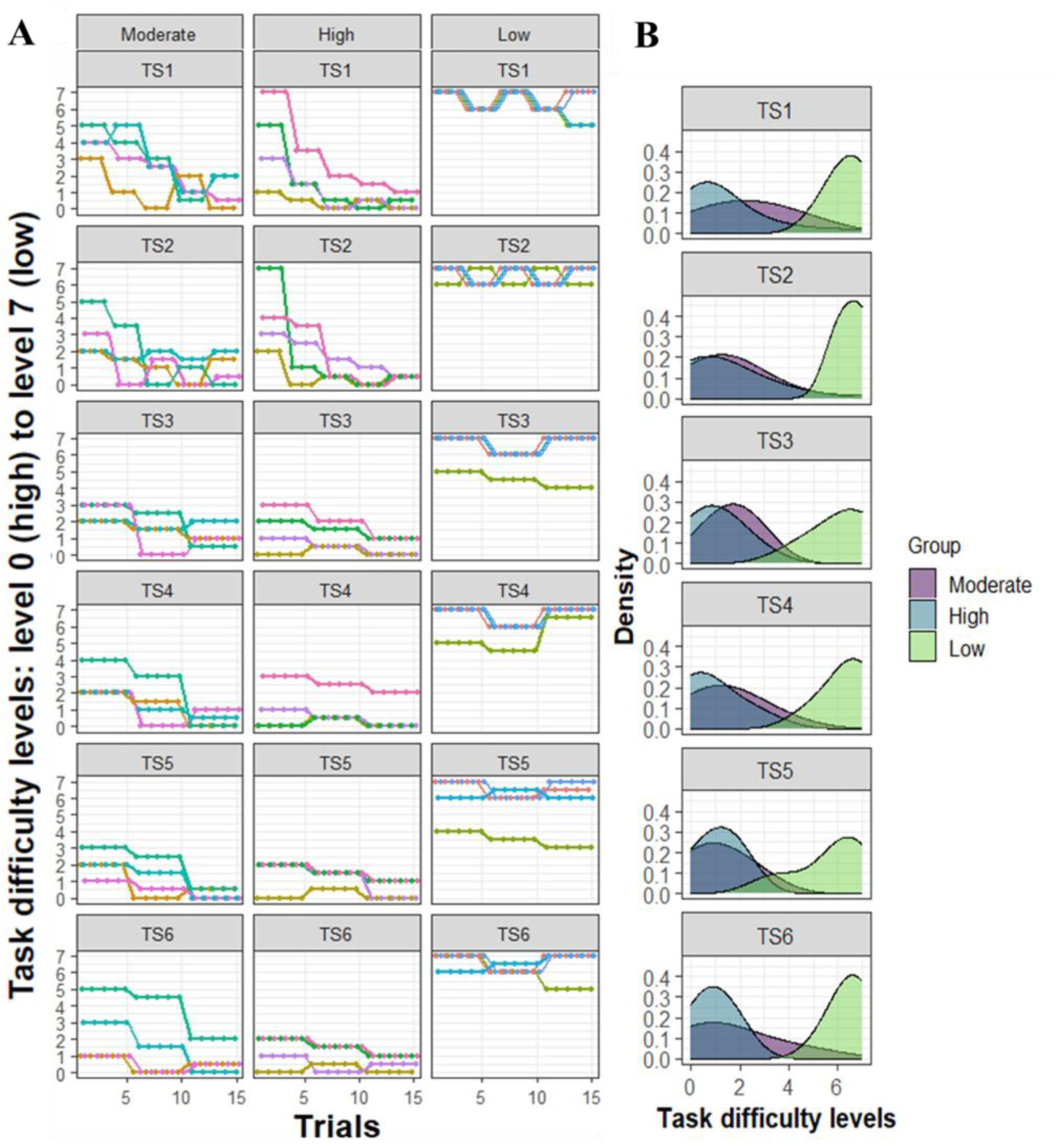
Manipulation of nominal task difficulty: (A) Participants were trained over 6 weeks where the DBT training load was manipulated based on their group assignment. Every training session was designed to start at an individualised load (task difficulty level) depending on the pretraining session DBT skill assessment. The training then progressed depending on their average TIB performance every three (TS1-TS2) or five trials (TS3-TS6). All participants experienced the same number of changes in level of task difficulty during each session to avoid potential contextual interference effects confounding the results (Magill and Hall, 1990). (B) Density plots show the distribution of the training load separately for each group at every training session. To illustrate the training regimes, the above figures show the training loads of four participants from each training group.

The training loads (level 0 – level 7) for all three groups were manipulated in order to maintain the functional task difficulty according to the assigned groups, i.e., moderate, high and low difficulty conditions. TIB exceeding the upper limit of the threshold by 1 sec meant increasing the task difficulty (by removing 1 plate), 2secs higher meant increasing the task difficulty by 1.5 plate and so on. Similarly, TIB 1 sec below the lower limit meant lowering the task difficulty (by adding 1 plate), and so on. If the participants remained within the threshold, the task difficulty increased by removing 0.5 plate (2.5kgs) for the next block. All participants experienced the same number of changes in level of task difficulty during each session (Contextual interference effect; Magill and Hall, 1990).

Fig. 3 shows an illustration of the maintained level of functional task difficulty (TIB) by manipulating the nominal task difficulty (from Fig.2) for four participants from each group.

**Figure 3.**
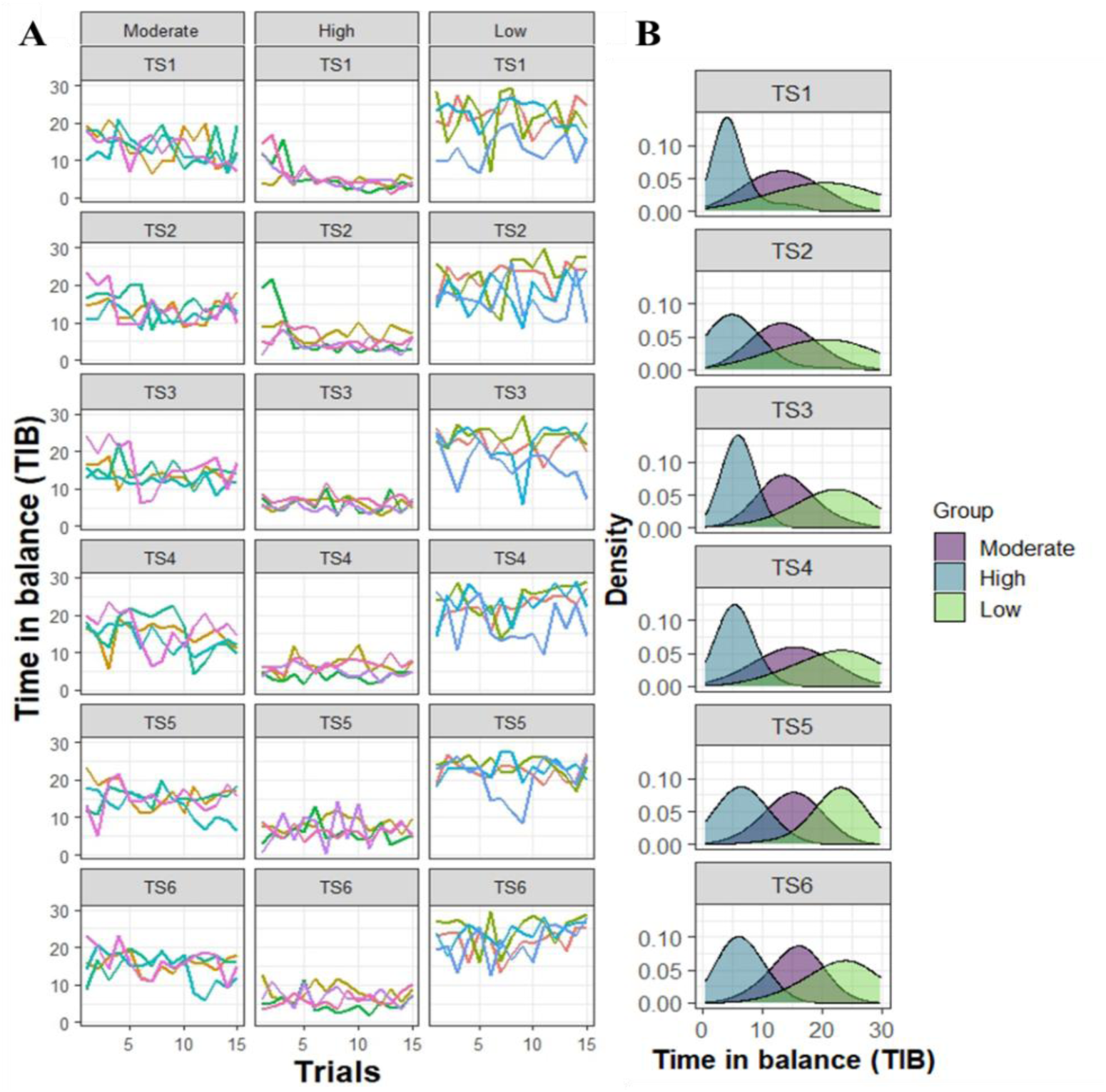
Maintenance of functional task difficulty: (A) Participants were trained on the DBT over 6 weeks where the training progressed with the aim of maintaining a predefined functional task difficulty (TIB) throughout each session. The threshold for progression depended on the group assignment, viz., the TIB threshold for high group: 3-6 secs; the moderate group: 12-15 secs; the low group: 23-26 secs. Training loads were manipulated (Fig. 2) based on these thresholds. (B) Density plots show the distribution of the maintained functional task difficulty (TIB) separately for each group at every training session. To illustrate the training regimes, the above figures show the DBT performance of four participants from each training group.

### 2.3. Transfer tasks (pre, inter/TS4 & post-test)

Transfer tasks were categorised into near motor transfer, far motor transfer, executive cognitive transfer and memory cognitive transfer tasks.

#### 2.3.1. Near motor transfer tasks (novel variations of DBT task)

- Added weight vest: In this task the participants performed the DBT task at the highest nominal task difficulty (level 0) while wearing a 6kg weight vest, thus, was considered as a near motor transfer task. A characteristic specific to the person was altered by artificially increasing the body weight via a weight vest while performing the task. This moves the COM higher and makes it heavier, hence making it more difficult to control. The variable of interest was the time in balance (TIB) from best of two trials of 20s each.
- Board position change: The DBT task was altered by changing a task-specific characteristic, i.e., using another stabilometer and moving the board further away from the rotating axis hence increasing the nominal difficulty of the task. Consequently, in this near motor transfer task, the movement dynamics were altered to make it more unstable while the participants performed a variation of the DBT at level 0. Similar to the above task, the variable of interest was the time in balance (TIB) from best of two trials of 20 s each.

#### 2.3.2. Far motor transfer

Wii task: An interactive football task available on a Nintendo WiiFit console was used to assess the participants’ ability to control their COM and goal-directed lateral sway (Goble et al., 2014). A Wii Nintendo connected to a large TV screen was used for this task. The participants stood with both feet on the Wii-board which tracked their centre of mass and represented it as an avatar on the screen. The goal of this task was to earn points by heading footballs kicked in their direction. Simultaneously, other objects (panda masks and shoes) were also randomly added to the mix with instructions to dodge these objects or risk losing points. These objects were tossed either laterally (left or right) or in the centre. This prompted the participant to accurately realign the avatar in the required direction by shifting their COM/ body weight while standing on the Wii board. Participants performed 10 trials at the beginners’ level with 30s breaks in between. The total score is derived based on the successful hits and misses of all objects during the trial. This score is indicative of the accuracy as well as reaction time of each participant.

#### 2.3.3. Executive transfer tasks

- D2- Test of Attention: This task assesses sustained attention, visual scanning speed and accuracy (Bates and Lemay, 2004). We used the German paper and pencil version of the revised “d2 Test of Attention” (d2-R; Brickenkamp R et al., 2010) consisting of 14 test lines with 47 symbols per line. The participant is required to strike through every occurrence of the letter “d” bearing only 2 dashes as quickly and accurately as possible. All other symbols act as distractors meant to be ignored. The symbols can be either lowercase letters “d” or “p” marked with 1, 2, 3, or 4 small dashes above and/or below the letter. The participants proceed from left to right with 20 seconds dedicated to each line; after which they start with the next line. Due to a low learning effect, the same version of the test was used for all three testing timepoints (pre, TS4, post-test). Concentration score was the variable of interest, defined as the number of marked distractors (sum of errors of commission and errors of overlooking) deducted from the total number of processed targets.
- Trail making test (TMT; Lezak et al., 2004; Rodewald et al., 2012): This task is considered as a measure of divided attention and scanning abilities, with particular focus on cognitive flexibility involving switching (between sets of letters and alphabets). This test included 2-parts: (i) part A required the participant to connect randomly scattered numbers starting from 1 to 25 in an ascending order; (ii) part B required the participant to alternate between letters (A-L) and numbers (1-13), i.e., connecting a number to an alphabet moving in an ascending order and alphabetical order respectively. Part B requires constantly switching between sets (1-A-2-B-3-C-…). The participant first is familiarised with a shortened version of each part before they begin. They are instructed to perform the task as fast as possible with minimal errors. Time to completion and difference between the time required to complete parts A and B (delta/Δ TMT (Corrigan and Hinkeldey, 1987; Drane et al., 2002)) are used as variables of interest. To avoid a learning effect, each subtest was mirrored horizontally or vertically for use during the intermediate (TS4) and post-test respectively.

#### 2.3.4. Memory transfer tasks

- Rey-Osterrieth Complex figure test (ROCF) and modified Taylor figure (Osterrieth, 1944; Rey, 1941; Tombaugh and Hubley, 1991): ROCF and modified Taylor figure (used as parallel version) are widely used to assess the visuo-constructional abilities and visuospatial recall memory, a non-verbal form of memory. Multiple cognitive domains are said to be involved in the execution of this test, viz., memory, attention, concentration, fine-motor coordination, visuospatial perception, planning, organization and spatial orientation; functions mediated by the prefrontal lobe, hippocampus and associated temporal regions (Shin et al., 2006; Zhang et al., 2021). This test consists of a copy trial phase requiring the participants to copy the complex figure placed in front of them as accurately as possible, followed immediately by the second step where they draw the figure from memory (immediate recall). The third phase is conducted 30 mins after the immediate recall phase where the participants are once again required to draw the figure from memory. The modified Taylor figure was used for the intermediate testing (TS4) and a mirrored version of the Rey Figure was used for the post-test. The variables of interests were the number of recalled items and the Rey quotient calculated as a percentage of recalled and copied items.
- Symbol Digit modalities test (SDMT; Boake, 2002; Sheridan et al., 2006): This test is a measure of associative learning, motor speed, attention, visuo-perceptual scanning ability and working memory (Jaeger, 2018). This task requires the participants to match as many symbols to numbers within 60 secs with the help of a key (set of pre-matched symbol-digit pairs given at the top of the sheet). The variable of interest was the number of correctly matched items. Parallel versions were used for the intermediate (TS4) and post-testing timepoints.
- Verbal learning and memory test (Helmstaedter, C., Lendt, M. & Lux, 2001): This task is a measure of declarative verbal memory, learning performance and long-term encoding-recall-retrieval performance (Spilkin et al., 2009). Participants are presented a set of 15 words verbally which are to be memorized and repeated out loud, irrespective of the order. This was performed 5 times, following which a set of 15 interfering words were presented; this time the participants had to call out as many interfering words as they could memorize. After the interference phase, the participants were asked to call out as many words as they could remember from the original list (previously repeated 5 times), this concluded the immediate recall phase. Thirty minutes after this phase, the delayed recall phase was conducted, where the participants were once more asked to call out loud the words from the original list. Parallel versions of this test were used for the intermediate (TS4) and post-testing timepoints. The variables of interest were the sum total words remembered after 5 repetitions of the original word list.

### 2.4 Data analysis

All statistical analyses for this study were conducted using the software R version 4.2.2 (R Core Team, 2022). Due to non-normal data distributions in the outcome measures, statistical analyses were performed using robust non-parametric statistical tests. Between group comparisons at baseline for all demographic variables were conducted using chi square or Kruskal-Wallis rank sum test (depending on the scale level).

Changes in balance (DBT) and transfer performance over time were analysed using RM-ASCA+ (Madssen et al., 2021), as implemented in R’s ALASCA package (Jarmund et al., 2022). RM-ASCA+ extends ANOVA–simultaneous component analysis (ASCA) to repeated-measures designs by combining linear mixed models (LMMs) with principal component analysis (PCA) (Madssen et al., 2021).

Here, the first step involved fitting LMMs separately for each task using the lme4 package (Bates et al., 2015). Prior to modelling, all variables were scaled by dividing each value by the standard deviation of the corresponding variable across all baseline (pre) samples. Time and the interaction between time and group were included as fixed effects, while participants (ID) were specified as a random intercept to account for within-subject correlations. Age, sex, and body weight were included as covariates. In line with recommendations for randomized controlled trials (Jarmund et al., 2022), the main effect of group was excluded (1). Reference coding was applied, with the moderate difficulty group and the pre-test time point serving as reference group/levels. The resulting LMM formula was as follows:

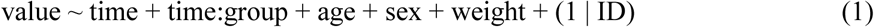

Complementary p-values for fixed effects in the LMMs were calculated using Satterthwaite’s method via the lmerTest package (Kuznetsova et al., 2017). To account for multiple testing across time points, the Benjamini–Hochberg procedure was applied, and adjusted p-values < .05 were considered statistically significant.

Fixed-effect matrices from these univariate LMMs were then subjected to PCA to derive principal component (PC) scores and loadings. PC1 is the line that best accounts for the shape of the points and represents the maximum variance direction in the data (Fig. 4 & 6). Loadings reflect the contribution of each task to temporal changes and group-by-time interactions; positive (or negative) loadings correspond to higher (or lower) values at time points associated with high PC scores.

**Figure 4:**
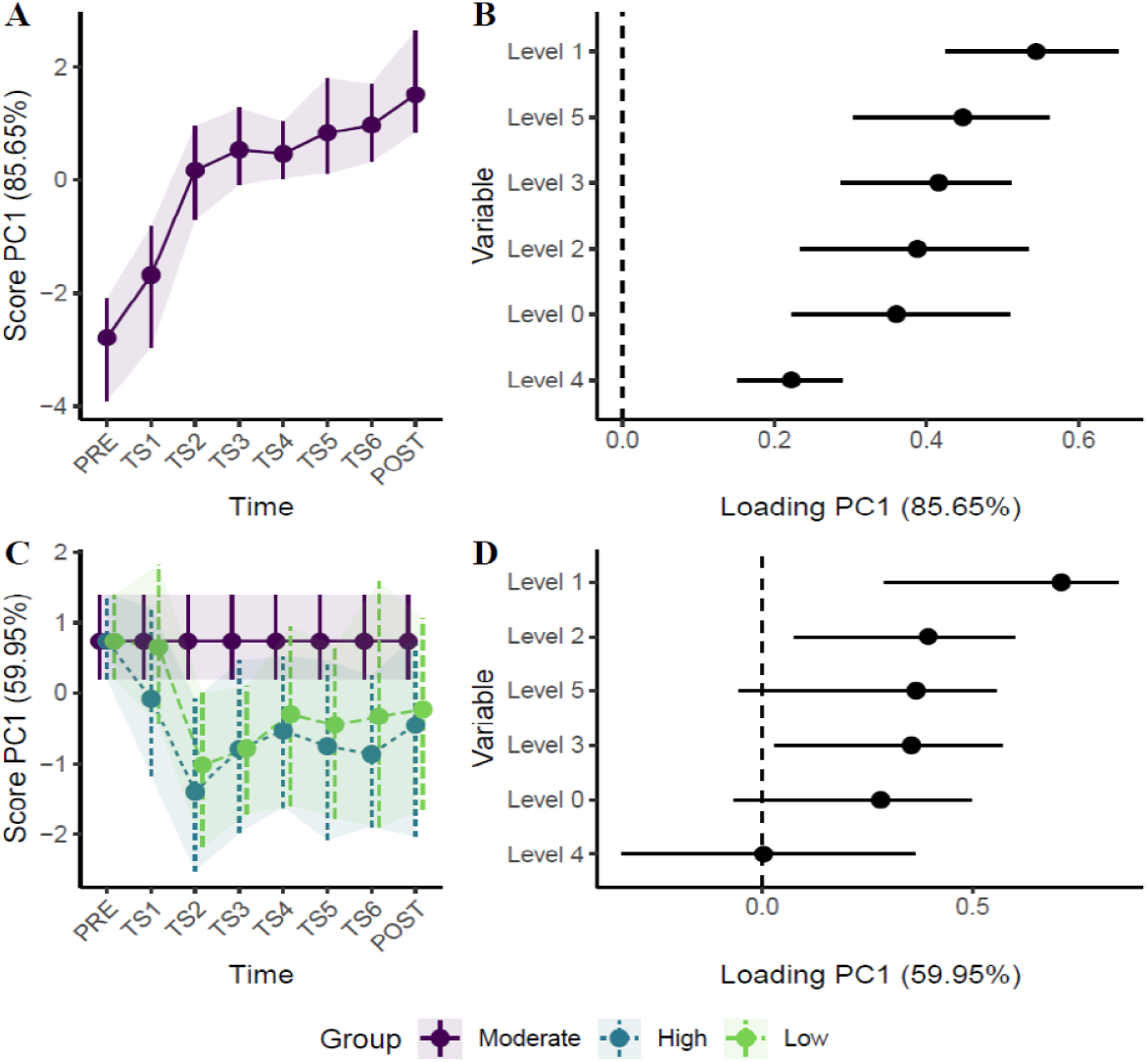
DBT performance improvements over time: (A) PC1 scores showing improvements in DBT performance of the moderate group (reference group) over the entire training duration; (B) factor loadings indicate a positive correlation between the different task difficulty levels (variables: level 0 to level 5) and DBT performance (PC1) of the moderate group; (C) between group differences in DBT improvements over the entire training duration. The time development of the moderate group has been removed to highlight the effect of training; (D) factor loadings indicate contributions of the different task difficulty levels (variables: level 0 to level 5) on the between-group differences in DBT performance (PC1).

To assess robustness, nonparametric bootstrapping (1,000 runs) were used to generate 95% confidence intervals (CIs) for scores and loadings. Statistically meaningful multivariate changes were inferred when CIs for PC1 scores did not overlap (Madssen et al., 2021). Likewise, loadings were deemed significant when their CIs excluded zero.

For the assessment of longitudinal changes in balance skill, participants were examined based on their performance (TIB/ value) on each task difficulty level (level 0 – level 5) separately at every timepoint. The transfer tasks were categorized into either near motor transfer, far motor transfer, executive cognitive transfer or memory transfer tasks (variables). For transfer task analyses, all raw scores were transformed into z-scores as the scales for each task differed. Z-scores (value) from each task were averaged over the respective categories.

Correlation analyses were conducted using the Spearman rank-based correlation for each group. PC1 scores obtained from the ALASCA analyses for DBT and transfer performances were used for these correlations. Lastly, the correlation coefficients for each group were aggregated using the Olkin-Pratt (OP) meta-analytical approach, treating the correlation coefficients as effect sizes. This was implemented via the metacor function in R (Laliberté, 2019; Schulze, 2004).

## 3. Results

Baseline characteristics of the participants in this study with respect to age, gender, body height, body weight, MoCA test and day-to-day physical activities did not differ between groups (Table 1).

**Table 1.** Demographic data: Comparisons between groups in relation to age, sex, height, weight, MoCA (cognitive assessment) and physical activity. Values displayed denote the median and interquartile ranges within parentheses for all groups. All statistical comparisons performed with Kruskal-Wallis rank sum test.

| Characteristics | Overall | High | Low | Medium | Between-group comparison |
| --- | --- | --- | --- | --- | --- |
| Age | 67 (5.75) | 67 (1.75) | 67.5 (7.75) | 67.5 (6.5) | H(2) = 0.014, $p = .99$ |
| Sex (F; M) | 18;12 | 6;4 | 5;5 | 7;3 | H(2) = 0.81, $p = .67$ |
| Weight | 72.08<br>(18.25) | 73 (13) | 75.85<br>(16.5) | 67.0<br>(22.29) | H(2) = 2.05, $p = .36$ |
| Height | 168.5 (12) | 167.5 (6) | 174 (10.75) | 166 (14.5) | H(2) = 0.61, $p = .74$ |
| MoCA | 26 (3) | 25.5 (4.75) | 26.5 (2.75) | 26.5 (3) | H(2) = 0.76, $p = .68$ |
| Physical activity level | 159.34<br>(107.66) | 185.9<br>(170.52) | 145.8<br>(34.69) | 202.03<br>(153.48) | H(2) = 2.18, $p = .34$ |

### 3.1. Effect of different training loads on balance ability

The DBT performance measured using the balance skill assessment conducted before every training session is also a measure of the retained DBT skill from the training conducted a week ago. The improvements in DBT performance of the moderate group (reference group) over the entire training duration is represented in Fig. 4A; here, PC1 explains 85.65% of variance in the main effect of time. Between-group differences are depicted in Fig. 4C, where PC1 explains 59.95% of between group variance (time x group interaction; the suboptimal groups: high and low vs the optimal/moderate group). Fig. 4C shows the divergence of the suboptimal groups from the reference group in terms of their DBT improvements. Non-overlapping CI’s at TS2 (Fig. 4C) indicate higher learning gains seen as early as after the first training session for the moderate (95% CI [0.20, 1.40]) compared to both the suboptimal groups (high: 95%CI [-2.53, -0.09] and low: 95%CI [-2.17, 0.002]). Differences in learning gains continued after the second training session between moderate and low condition (TS3: 95%CI [-1.72, 0.08]). At TS6 on the other hand, a difference in learning gains was observed between the moderate and high group (95%CI [-1.90, 0.24]). Factor loadings displayed in Fig. 4B and 4D indicate a positive correlation between the different task difficulties within the balance skill assessment (variables: level 0 to level 5) and DBT performance (PC1). Large (either positive or negative) loadings indicate a strong effect of a variable on the principal component; placed in the order of influence from the highest (at the top) to the lowest (at the bottom). Higher factor loadings at level 1, 2, 3 and 5 indicate that the improvements of the moderate group and the between group differences were strongly driven by these variables.

To further explore the DBT performance gains at each difficulty level, we looked at univariate results based on LMMs, with multiple testing correction applied via the Benjamini-Hochberg (1995) procedure (Fig. 5). Following a single training session (TS2), the moderate group exhibited significantly greater improvements at difficulty levels 1(high: *b* = -1.185, *p* < .05), 2 (high: *b* = -1.155, *p* < .05) & 5 (high: *b* = -1.263, *p* < .05) compared to the high group. Likewise, a significant difference was also found between the moderate and low groups at level 1 (*b* = -1.415, *p* < .05). Additional between-group differences in later training sessions, specifically at level 1 (TS5) and level 0 (post) were observed, the moderate group tended to outperform both the high group (level 1: *b* = -1.389, p < .05) and the low group (level 1: *b* = - 1.268, *p* = .07; level 0: *b* = -1.073, *p* = .09). Overall, the advantage of moderate (optimal) training was more pronounced at the early phases of learning and at higher task difficulty levels.

**Figure 5:**
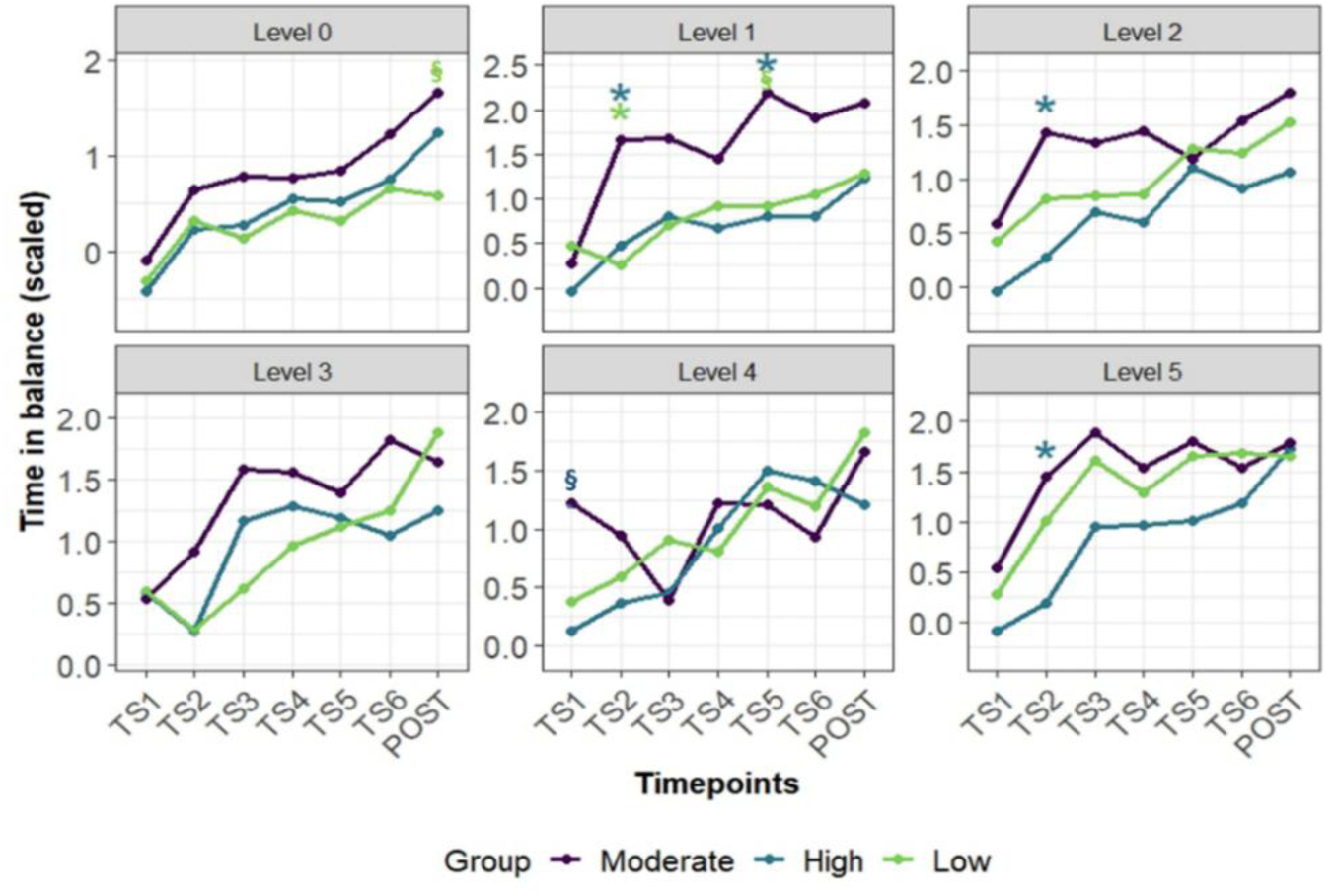
DBT improvements at each task difficulty level: The advantage of moderate training is observed at the early phases of training and at higher task difficulty levels. Significant differences (p < .05, FWE-corrected) are indicated using “*”, whereas trend towards significance (.05< p < .1, FWE-corrected) are indicated using “§”. Note that the range of the scales on the Y-axes for the difficulty levels differ due to the magnitude of improvements.

### 3.2. Effect of different training conditions on motor and cognitive transfer

Improvements in transfer task performance of the moderate group (reference group) is represented in Fig. 6A with 99.06% of variance explained by PC1(time effect); while the between group differences between the suboptimal groups and the moderate (optimal) group is displayed in Fig. 6C with PC1 explaining 93.23% of variance (time x group interaction). At TS4, differences in transfer performances were only detected between moderate (95% CI [0.32, 1.06]) and high groups (95% CI [-1.57, -0.26]), while at post-test, differences between moderate (95% CI [0.32, 1.06]) and both high (95% CI [-2.03, - 0.28]) and low groups (95% CI [-1.53, -0.06]) were observed. Loadings displayed in Fig. 6B and 6D indicate a correlation between the near motor, far motor, executive and memory transfer tasks (variables) and the PC1 scores. Near motor transfer tasks had a significantly higher contribution to the observed improvements in transfer performances of the moderate group and between group differences at TS4 and post-test.

**Figure 6:**
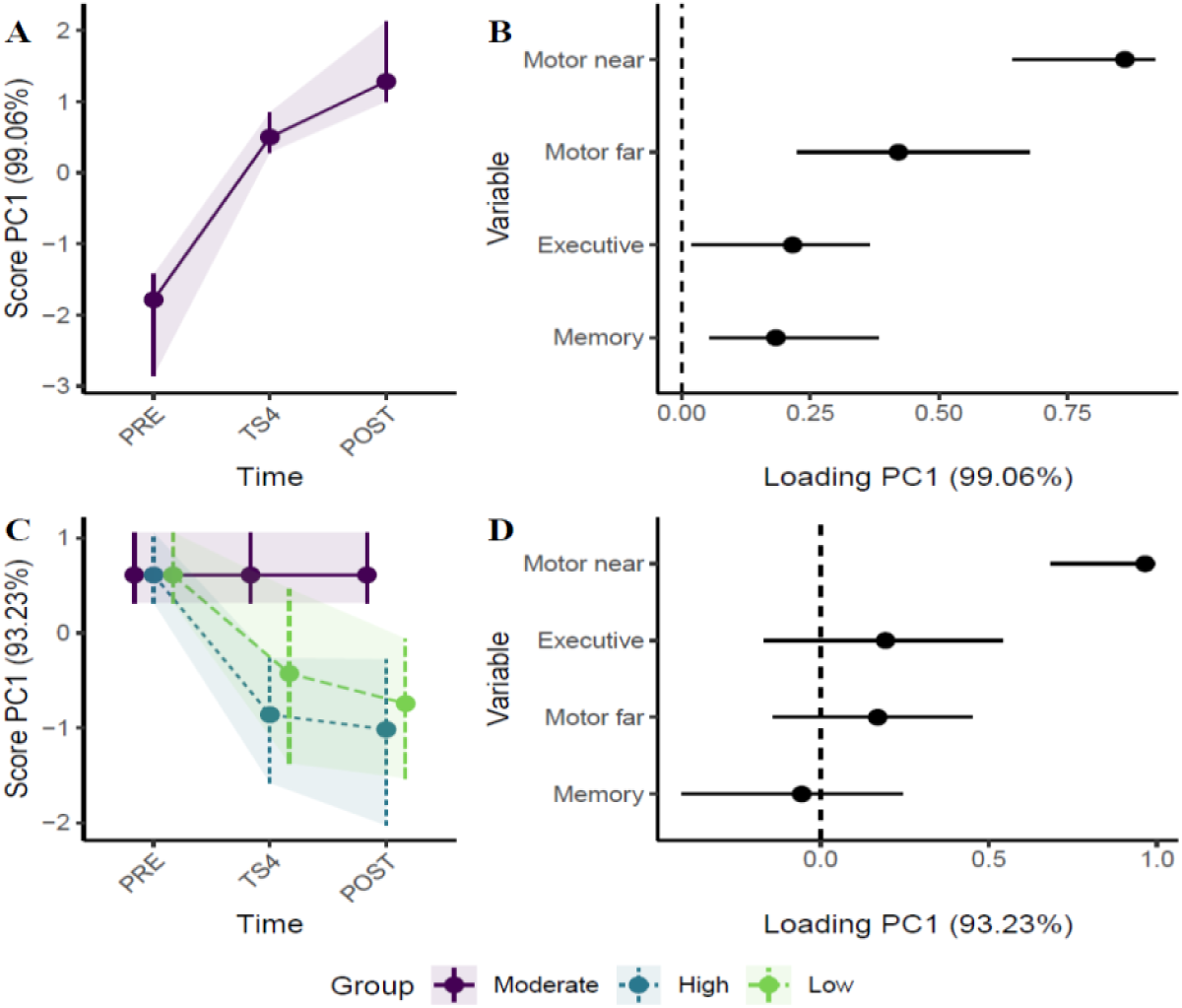
Motor and cognitive transfer tasks improvements over time: (A) PC1 scores showing improvements in transfer performance of the moderate group (reference group) over the entire training duration; (B) factor loadings indicating a positive correlation between the different transfer tasks (variable) and transfer performance (PC1) of the moderate group; (C) between group differences in transfer improvements over the entire training duration. The time development of the moderate group has been removed to highlight the effect of training; (D) factor loadings indicate contributions of the different transfer tasks (variables) on the between group differences in overall transfer performance (PC1).

Post hoc analyses of the observed transfer improvements (Fig. 7) revealed significant benefit of moderate difficulty training compared to both low (*b* = -1.354, *p* < .05) and high (*b* = -1.577, *p* < .01) training over near motor transfer tasks at post-test. Similar differences were already visible between the moderate and high groups (*b* = -1.361, *p* < .05) as early as after three training sessions (TS4). Moderate training load also influenced the executive transfer performance differences compared to the high group at TS4, however, this difference did not reach statistical significance (*b* = -0.478, *p* = .2).

**Figure 7:**
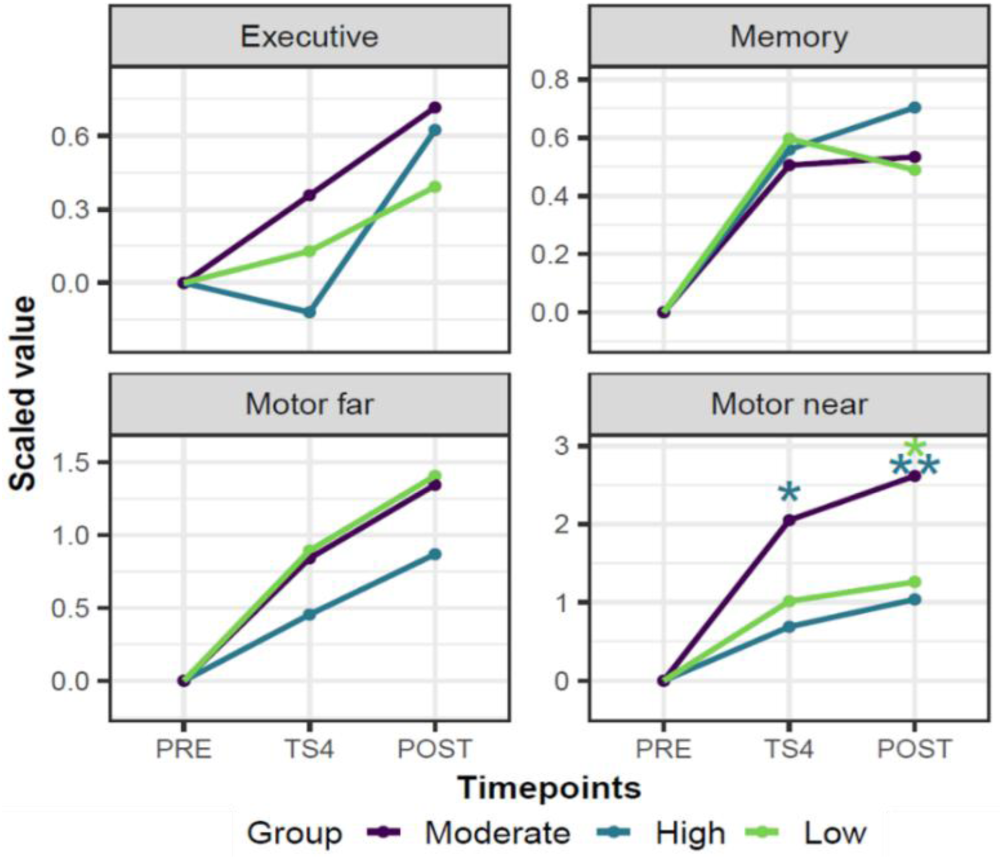
Performance improvements in each transfer task: Higher near motor transfer is observed as a result of moderate training at both TS4 and post-test. Significant differences (p < .05, FWE-corrected) are indicated using “*”. Note that the range of the scales on the Y-axes for all 4 tasks differ due to the magnitude of improvements.

### 3.3. Effect of DBT performance improvements on transfer

Overall changes in performance were observed for all groups after DBT training in terms of both balance performance and transfer. Therefore, correlational analyses on PC1 scores extracted from RM-ALASCA were performed to inspect if the observed transfer effects were related to DBT improvements. At TS4, the improvements in the transfer performance were positively correlated with the DBT improvements for the moderate group, Spearman *r* = 0.50 (moderate effect size), *p* = .14, the high group, Spearman *r* = 0.94 (strong), *p* < .001, and the low group, Spearman *r* = 0.56 (moderate), *p* = .09.

At post-test, similarly, the transfer performance positively correlated with the DBT improvements for all groups, viz., moderate group, Spearman *r* = 0.63 (moderate), *p* = .05, high group, Spearman *r* = 0.81 (strong), *p* = .008 and the low group, Spearman *r* = 0.29 (weak), *p* = .41.

Aggregating these correlations across using the Olkin and Pratt approach (Laliberté, 2019; Schulze, 2004), revealed effect size *G* = 0.69 (moderate), *p* < .0001 at TS4 and effect size *G* = 0.60 (moderate), *p* < .0001 at post-test. These results suggest that transfer effects were positively correlated with improvements in DBT performance irrespective of the training load.

## 4. Discussion

In this study, we manipulated DBT/ balance learning stimulus for older adults by matching the nominal balance task difficulty to their individual balance ability. This approach aimed to not only optimise motor learning but also augment transfer to untrained motor and cognitive tasks. The results showed early improvements in DBT performance after training under moderately difficult condition compared to suboptimal conditions (high and low difficulty training). These effects were more pronounced under challenging DBT test conditions with high task demands. Notably, near motor transfer effects were observed in the moderate group as early as after three training sessions (TS4/inter-test) which persisted post-intervention. No group differences were observed in far-motor and cognitive tasks. The results support predictions of the challenge point framework (Guadagnoli and Lee, 2004) and highlight the need for personalised intervention strategies for optimal motor learning.

### 4.1. Response-optimised training enhances task learning

Applying the Challenge Point Framework (CPF - Guadagnoli and Lee, 2004), we individualised DBT training by adapting *nominal* task difficulty during each practice session using a novel weight manipulation strategy. This strategy allowed us to experimentally apply distinct *functional* task difficulty levels in three different load conditions (Fig. 3). We tested predictions of the CPF using high, low and moderate (optimal) training load groups based on performance ranges from previous studies using the DBT (Aye et al., 2025; Lehmann et al., 2020; Sehm et al., 2014; Taubert et al., 2010). While all groups improved DBT performance, the moderate training group showed greater DBT improvements in the early period of learning. Similar results were obtained by (Akizuki and Ohashi, 2015), where they observed ineffective motor learning at the most and least difficult levels of task difficulty in young adults. Our findings extend these findings to motor learning in older adults. It is speculated that the early improvement in learning occurred as a result of enhanced task encoding through optimal load conditions (Lustig et al., 2009; Lustig and Flegal, 2008). The suboptimal groups, however, required additional sessions for similar gains. This early performance advantage achieved through moderate functional task difficulty levels ceased in the later learning period suggesting meaningful but transient skill-specific early benefits.

Following findings of dissimilar DBT gains between healthy and Parkinson’s participants (Sehm et al., 2014), our results underscore the role of individualised task demands in optimising motor learning trajectories in older individuals. Previous studies (e.g., Bootsma et al., 2020) have shown high task difficulty to impair learning and neural engagement, a phenomenon often observed in older adults (Lustig et al., 2009; Park and Reuter-Lorenz, 2009). Therefore, optimising task load can be used as a strategy to avoid task disengagement through maintenance of motivation by balancing challenge and success (Hodges and Lohse, 2022; Wulf and Lewthwaite, 2016).

Given age-related neural variability (Cabeza et al., 2018; Iordan et al., 2021), optimising task difficulty may support more efficient information processing creating an opportunity to maximise learning benefits. We believe, the moderate difficulty training (optimal stimulus in our study), was able to aid efficient information processing which is especially relevant in the early learning phase while learning a novel complex task (Guadagnoli and Lee, 2004). Furthermore, our data suggests early DBT gains in the moderate group were most pronounced under challenging DBT task demands, while suboptimal groups trailed at these levels (Fig. 5). More specifically, the moderate group outperformed the high group under challenging test conditions although both groups often experienced overlapping nominal task difficulties during training (Fig. 2). Interestingly, despite the low levels of nominal task difficulty experienced by the low group during training (Fig. 2) their DBT improvements under challenging task demands were often similar to that of the high group (Fig. 4 and 5). These findings are consistent with CPF and related studies that have shown low difficulty training loads to also be suitable for performance improvements in novice young and older adults (Al-Fawakhiri et al., 2023; Bootsma et al., 2021, 2020; Guadagnoli and Lee, 2004; Hodges and Lohse, 2022; Wilson et al., 2019). We interpret these findings as evidence for the beneficial effect of maintaining low-to-moderate levels of functional task difficulty on skill acquisition in healthy older adults. This advantage of the moderate group in DBT performance compared to low- and high groups (Fig. 4) is consistent with predictions of the CPF and suggest beneficial effects at approx. 40-50% of maximum DBT performance (error rate).

### 4.2. Response-optimised training promotes transfer

Beyond DBT improvements, optimally trained participants also showed enhanced performance in near-motor transfer tasks (Fig. 7), particularly those tasks that share characteristics with the trained DBT task. Performance in DBT with additional weight vest and with a larger distance between board pivot and its support surface was higher in the moderate group compared to both suboptimal groups. Typically, complex tasks like DBT requires managing multiple degrees of freedom and can foster a compilation of strategies (generalisable and specific) when practiced under discovery-learning conditions (Orrell et al., 2006). These strategies can help individuals transfer learned principles to new tasks with similar goals (Taatgen, 2021). The strategies developed on our trained task were easily applicable/generalisable onto untrained tasks that shared similar biomechanical characteristics, postural sway tactics, task goals, etc, despite being characteristically of a much higher nominal difficulty. Our results suggest encouraging strategy development over prescription enables self-generated processes which are easier to incorporate in unfamiliar contexts (Lustig et al., 2009; Lustig and Flegal, 2008). This approach combined with an optimally paced training likely equipped the moderate group to better adapt to closely related motor tasks throughout the training period (TS4 and post).

Successfully executing a task depends on its complexity and the experience the individual has with the task. In this respect, older adults in particular face high degree of challenge in terms of cognitive load while executing novel complex tasks (Cabeza et al., 2018; Guadagnoli and Lee, 2004; Tomporowski and Pesce, 2019). A tailored training with moderate levels of difficulty paced to the skill level of the individual, as applied here, enabled performance improvements through efficient information processing without overburdening their current capacities (unlike the high group) or disengagement (Wulf and Lewthwaite, 2016). Moreover, task variability stemming from gradually increasing nominal difficulty may have reduced specificity to the trained task, instead supporting generalisation to related tasks (Hodges and Lohse, 2022; Ranganathan et al., 2020).

Transfer improvements to more distant static postural tasks (e.g., Wii balance board) were similar across all three groups, however, less pronounced as compared to near transfer task improvements. As mentioned earlier, this may result from differences in the task characteristics between the Wii task and the trained task (static vs dynamic balance task). Although both tasks qualify as balance-related activities, the strategies utilised in both these tasks are not similar enough for participants to utilize the learned strategies (generalisable) in this new context (Giboin et al., 2015; Kümmel et al., 2016).

Even though cognitive transfer was hypothesised due to overlapping cognitive demands and previously observed commonalities in brain plasticity and brain stimulation studies between DBT and cognitive functions (Prabhu et al., 2024; Taubert et al., 2010), all groups improved comparably on executive and memory tasks. This suggests that distinct levels of functional DBT task difficulty had no differential effect on cognitive functions. While DBT training could have potentially challenged cognitive circuits in the brains of older adults, it seems that training load does not mediate cognitive transfer effects. Compared to healthy young adults, the impact of balance training on older adults’ frontal brain function seems to differ (Lehmann et al., 2025). Future studies are essential to test balance training characteristics and behavioural tests through which cognitive transfer is observable (Dordevic et al., 2017; Rogge et al., 2017).

### 4.3. Transfer performance correlates with DBT skill improvements

Transfer performance in general was positively linked to DBT gains in all groups (Fig. 8). Those with greater DBT improvements showed higher transfer, indicating that skill acquisition during training reinforces success in related tasks (Hodges and Lohse, 2022). Although the moderate group achieved higher overall performance improvements, the observed improvements in DBT were reliable predictors of transfer, regardless of training load and particularly for motor transfer tasks.

**Figure 8:**
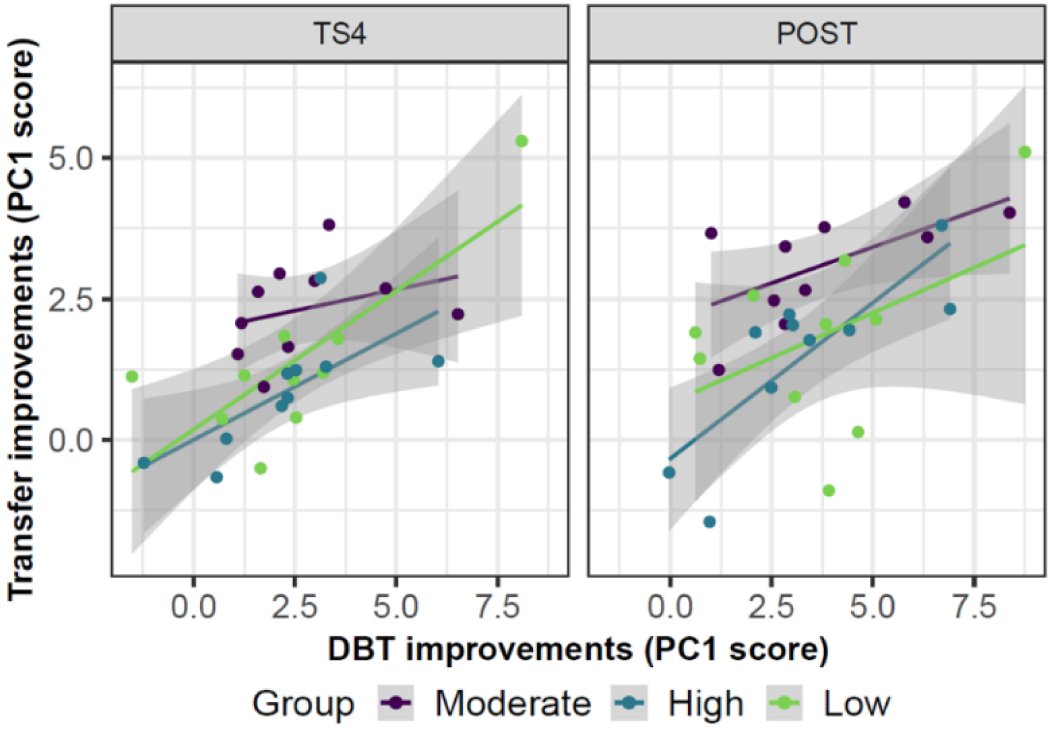
Correlation between DBT improvements and transfer task performances at TS4 and post-test.

Overall, optimised training conditions enhanced both immediate learning and transfer, underscoring the importance of individualised task difficulty in promoting motor adaptability and functional independence in older adults.

## 5. Conclusion

The results of this study pertaining to balance learning and near motor transfer support the theoretical concepts put forth by the challenge point framework (Guadagnoli and Lee, 2004). To the best of our knowledge, this is the first study to implement the CPF in a long-term motor training paradigm in older adults. Optimisation of task difficulty accounting for the individual’s abilities and the surrounding practice variables may lay the foundation for efficient learning by shortening training duration. In particular, the concept of individualisation of training loads along with dynamic alteration of functional task difficulties (response-optimisation) are key aspects to be implemented in future studies looking to further validate the CPF and some of its concepts not tested in this study (random-practice, modelled information in practice, varying frequency of knowledge of results, etc). The affordance of a higher error rate (∼ 40-50%) as the optimal challenge point in our study is probably due to a longer training duration as compared to previous single training session studies that suggest a lower error rate (Al-Fawakhiri et al., 2023; Bootsma et al., 2021, 2020, 2018; Wilson et al., 2019). In addition, a potential task-specific dependency of the optimal challenge point needs to be explored. The transfer effects, however, were not entirely dictated by the training loads. Instead our results show that individuals benefiting from DBT training in turn may also be able to capitalise on vital skills learned during training to improve performance on related tasks.

In conclusion, our findings demonstrate that the benefits of response-optimisation of practice conditions in old age are not only limited to greater learning gains on the trained task, but also reflect performance improvements in closely related motor tasks. These results further highlight the potential for transfer and the training conditions that enable it, hence, encouraging exploration of its underlying functional mechanisms. Through this study we were able to harness the existing strengths of older individuals to further build capacities in the face of progressive age-related physical and cognitive deterioration.

## Data availability

Data will be made available upon requests addressed to corresponding authors (NP or MT).

## Acknowledgements

The authors would like to thank Johannes Riedel, Henriette Kramer and Christoph Klötzer for their support in data acquisition for this study. We would also like to acknowledge assistance provided by Naemi Ristl, Sophie Lübke and Kolya Laade for piloting various methodological aspects in preparation of this study.

## Author contributions (CRediT)

NP: Conceptualisation, methodology, data acquisition, data curation, formal analyses and writing-original draft, review and editing;

MM: Conceptualisation, methodology, data curation and writing-review and editing;

NL: Conceptualization, methodology, supervision, formal analyses and writing-review and editing;

NA: Conceptualisation, methodology and writing-review and editing;

GZ: Conceptualization, methodology, resources, funding acquisition, writing-review and editing;

MT: Conceptualization, methodology, resources, funding acquisition, supervision and writing-original draft, review and editing.

## Author statement

We declare that this manuscript is original, has not been published before and is not currently being considered for publication elsewhere. We confirm that the manuscript has been read and approved by all named authors and that there are no other persons who satisfied the criteria for authorship but are not listed. Generative AI or AI-assisted technologies were not used at any point in this work.

